# Epithelial NCAPD3 expression protects against stress-induced intestinal injury in mice

**DOI:** 10.64898/2026.04.21.719792

**Authors:** Isabel Johnston, Erin E. Johnson, Afshin Khan, Michelle S. Longworth, Christine McDonald

**Author notes:** Corresponding author (CM).

## Abstract

Intestinal epithelial cells are central players in mucosal barrier integrity and host-microbe interactions. Genetic studies have revealed that epithelial dysfunction is a key contributor to the pathogenesis of inflammatory bowel disease. Non-SMC condensin II complex subunit D3 (NCAPD3) is essential for chromatin organization and stability. NCAPD3 also promotes antimicrobial defense and autophagy responses *in vitro*. NCAPD3 expression is decreased in intestinal epithelial cells from patients with ulcerative colitis; however, it is not known whether loss of NCAPD3 expression drives intestinal barrier dysfunction or is a result of disease-associated inflammation. To investigate this relationship *in vivo*, a tissue-specific approach was required, as global constitutive knockout of NCAPD3 is embryonic lethal. Therefore, a transgenic mouse line with doxycycline-inducible expression of a short hairpin RNA targeting NCAPD3 restricted to villin-expressing cells was generated (NCAPD3KD mice) to enable the study of NCAPD3 function in the intestinal epithelium. Treatment of NCAPD3KD mice with 9-tert-butyl doxycycline resulted in ∼75% reduction of NCAPD3 protein in EpCAM_+_ intestinal cells. Short-term epithelial NCAPD3 knockdown did not induce spontaneous colitis but was associated with increased serum amyloid A and a trend towards increased intestinal permeability. Upon dextran sodium sulfate or *Salmonella enterica* serovar Typhimurium ΔAroA challenge, NCAPD3KD mice exhibited exacerbated weight loss, higher disease activity, increased histopathological damage, abnormal colonic cytokines and chemokines, and significantly increased intestinal permeability. These results indicate that NCAPD3 expression in the intestinal epithelium is required for optimal barrier maintenance and antimicrobial defense under chemical or microbial stress. These findings support prior *in vitro* observations and solidify NCAPD3 as a regulator of intestinal epithelial barrier function and mucosal host defense.

**Author Summary:** NCAPD3 is a multifunctional protein with established roles in chromatin organization, genome stability, mitochondrial function, and antimicrobial defense. Dysregulated NCAPD3 is implicated in human diseases, such as inflammatory bowel disease (IBD) and microcephaly; however, due to its essential role in cellular division, determination of whether NCAPD3 loss drives these pathologies *in vivo* has been lacking. Using a new transgenic mouse model that selectively reduces NCAPD3 expression in intestinal epithelial cells, our study establishes NCAPD3 as an epithelial regulator of the mammalian intestine that enhances epithelial barrier resilience and antimicrobial defense during stress. Although dispensable for short-term basal homeostasis, NCAPD3 function becomes critical during epithelial injury and enteric infection. Reduced NCAPD3 expression may therefore lower the threshold for inflammatory disease by weakening barrier integrity, amplifying inflammatory cascades, and impairing antimicrobial defenses. These findings position NCAPD3 as a potential modulator of IBD susceptibility and highlight chromatin organization as an important, previously underappreciated layer of intestinal epithelial regulation.

## Introduction

The intestinal epithelium is a vital single-cell barrier between the host and the external environment, consisting of a diverse range of specialized intestinal epithelial cells (IECs) (1). This dynamic interface allows the absorption of nutrients and fluids, provides innate immune defenses, and helps maintain a homeostatic balance with the gut microbiota (2). The epithelium provides a physical barrier, with intercellular tight junctions responsible for maintaining a seal between adjacent cells (3). Additionally, IECs convey signals to mucosal immune cells and help coordinate differential immune responses to commensal and pathogenic bacteria. Defects in epithelial barrier integrity are a hallmark of inflammatory bowel diseases (IBD), such as ulcerative colitis (UC) and Crohn’s disease (CD), and render the host susceptible to enteric infections (2).

In the context of IBD, disruption and dysregulation of IEC function are key drivers of chronic intestinal inflammation (4). Defects in epithelial barrier integrity, including altered tight junction architecture, impaired mucus production, and reduced antimicrobial peptide secretion, lead to increased intestinal permeability and inappropriate exposure of the mucosal immune system to luminal microbes and antigens. Increased exposure of the host tissues to microbial products, from a so-called “leaky gut”, leads to chronic inflammation. Beyond barrier failure, IECs in IBD display aberrant innate immune signaling, stress responses, and altered interactions with the microbiota, all of which contribute to sustained inflammatory responses characteristic of IBD. Importantly, IBD has a clear genetic component, with genome-wide association studies identifying over 250 risk loci, many of which are directly linked to epithelial cell function (5). Collectively, these findings highlight IECs as central regulators of intestinal homeostasis and underscore how IEC dysfunction can predispose individuals to IBD.

Non-SMC condensin II complex subunit D3 (NCAPD3) is a core component of the condensin II complex, which promotes chromatin condensation during mitotic initiation (6) and maintains genomic stability (7). Beyond its canonical nuclear functions, NCAPD3 has been implicated in retrotransposon silencing (8), mitochondrial redox (9), and antimicrobial defenses (10, 11). Consequently, the dysregulation of NCAPD3 has been associated with multiple human pathologies. Loss-of-function mutations drive microcephaly (12), while overexpression has been associated with cancer progression (13-19). Critically, reduced NCAPD3 expression has been observed in intestinal epithelial cells isolated from UC patients (11). However, despite evidence from cell-based and *ex vivo* analyses of intestinal epithelial cells, the role of NCAPD3 in maintaining intestinal epithelial homeostasis *in vivo* remains uncharacterized. Specifically, it is unclear how loss or dysregulation of NCAPD3 affects epithelial integrity or susceptibility to intestinal inflammation within an organismal context.

To directly define the role of NCAPD3 in the intestinal epithelium *in vivo*, we created an inducible, intestinal epithelial cell-specific NCAPD3 knockdown mouse model. Using this model, we examined whether epithelial NCAPD3 is necessary to maintain intestinal homeostasis or to protect the intestinal barrier during inflammatory or infectious challenges. Specifically, we assessed the effects of NCAPD3 depletion on intestinal permeability, susceptibility to chemically induced epithelial injury, and host defense against enteric bacterial infection. Our studies provide evidence that NCAPD3 expression in the intestinal epithelium is essential for optimal barrier integrity and antimicrobial defense under stress from chemical or microbial challenges. These findings underscore the important role of NCAPD3 in maintaining intestinal health.

## Results

### Development of an intestinal epithelial cell-specific inducible NCAPD3 knockdown murine line

To investigate the role of NCAPD3 in the murine intestinal epithelium, we designed mice with inducible expression of a shRNA targeting NCAPD3 restricted to villin-expressing cells. Candidate shRNA sequences were screened for NCAPD3 knockdown efficiency prior to the creation of the knock-in transgene cassette (**Fig. S1A & Table S1**). The shRNA sequence targeting the 3’ untranslated region that reproducibly reduced NCAPD3 levels over 50% was used for the knock-in cassette. A CRISPR/Cas9-mediated targeted integration strategy of the transgene cassette into the *Col1a1* safe harbor locus was used to minimize the potential impact of the knock-in on the expression of other genes (**Fig. 1A**). Three injection conditions with different ratios of Cas9/small guide RNA/targeting vector were used to create targeted B6SJL/F2 fertilized oocytes that were then transferred to CD1 pseudo-pregnant recipients. The resulting 78 pups (founders) were then screened by qPCR and confirmed by sequencing of PCR products for transgene insertion and orientation (**Fig. S1B**). Transgene insertion was detected in 25.6% of the founders, and 85% of the transgenes were inserted in the correct orientation (21.8% overall). Of the mice with correctly oriented transgenes, germline transmission occurred in 40% of the lines.

**Figure 1:**
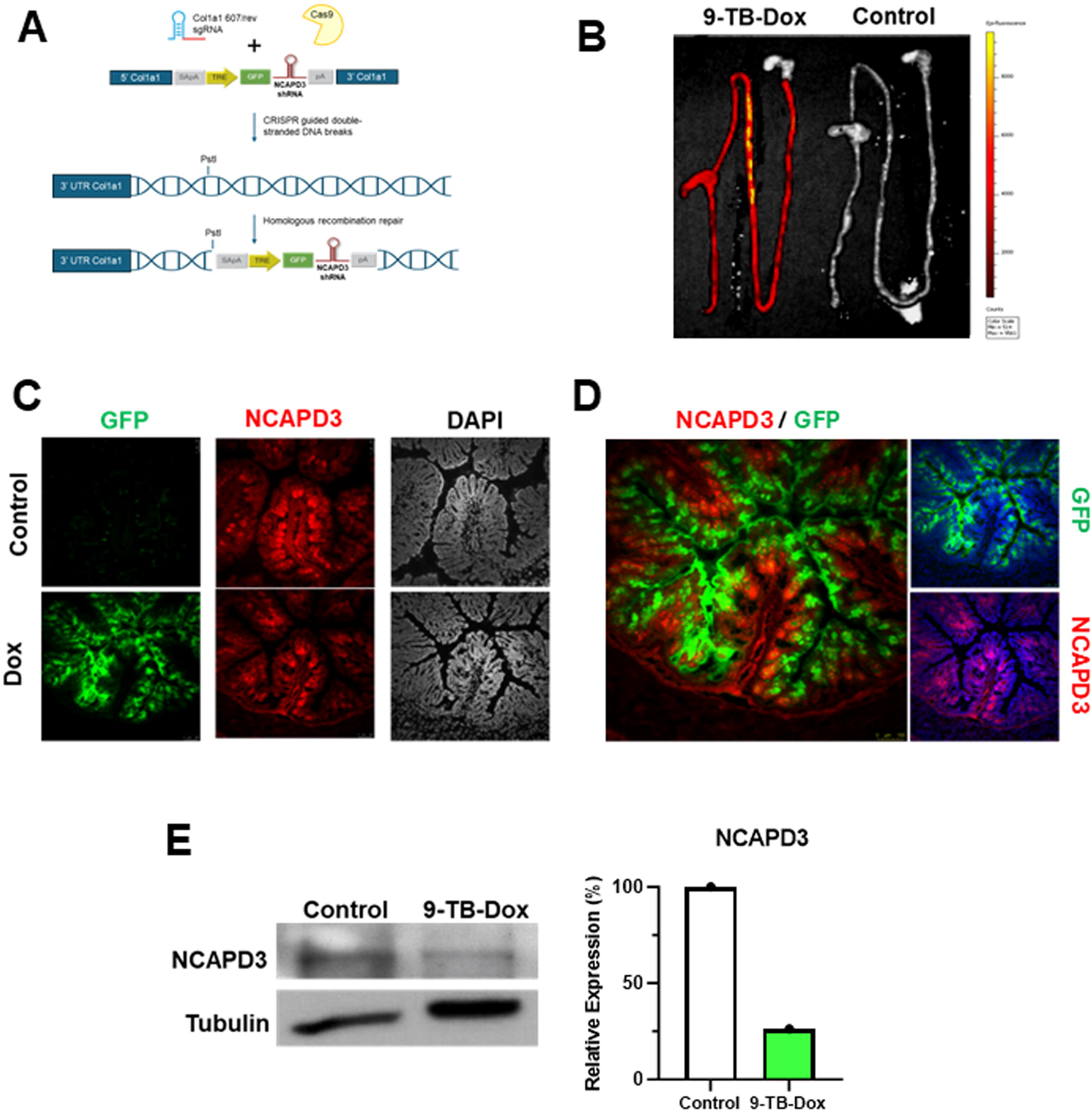
Design and characterization of NCAPD3KD mice. **A**. Schematic of transgenic targeting cassette and genomic insertion strategy. **B**. Intestinal expression of transgene cassette confirmed by fluorescent imaging of intestinal tissues *ex vivo*. Mice were injected with saline (Control) or 9-TB-Dox (25 mg/kg ip) for 4 days. Tissues were excised and imaged on an IVIS small animal imager. **C**. Visualization of NCAPD3 knockdown in colonic tissue by immunofluorescent confocal microscopy. Mice were treated with 9-TB-Dox (25 mg/kg; i.p. daily) for 2 days, and intestinal tissues were excised. Induction of transgene cassette visualized by detection of GFP reporter gene (green), NCAPD3 (red), and nuclei stained with DAPI (grey). Slides imaged with 10x objective. **D**. Larger image presentation of NCAPD3 (red) and GFP (green) colonic expression of 9-TB-Dox treated mice from C to visualize the non-overlapping signals of NCAPD3 and GFP in the colon epithelium. Nuclei stained with DAPI (blue). **E**. Quantification of intestinal epithelial NCAPD3 levels by immunoblot. Intestinal epithelial cells were isolated from mice treated with 9-TB-Dox (25 mg/kg i.p.) for 4 days, and Epcam^+^ /CD45^-^ cells were sorted via flow cytometry for GFP positivity. NCAPD3 protein levels were assessed by immunoblotting relative to a tubulin loading control and quantified using ImageJ.

Induction of transgene cassette expression and knockdown of NCAPD3 protein in intestinal epithelial cells in response to a non-antimicrobial tetracycline analogue, 9-TB-Dox (20), was evaluated by fluorescent imaging, immunostaining, and immunoblot analyses. As the microbiota contributes to intestinal health, 9-TB-Dox was used to activate expression of the NCAPD3-targeting shRNA and GFP reporter gene instead of the antibiotic tetracycline or doxycycline to minimize the impact of the inducing agent on the microbiota. After 4 days of daily 9-TB-Dox i.p. administration, gross evaluation of tissues by fluorescent imaging confirmed selective expression of the GFP reporter gene throughout the small intestine, cecum, and colon (**Fig. 1B**). Examination of extraintestinal tissues also known to express villin, such as the pancreas and kidney, had low level GFP expression; however, fluorescence levels in these tissues were approximately 50-fold lower than those observed in the intestine.

Closer evaluation of intestinal tissue by confocal fluorescent microscopy showed epithelial-specific GFP expression after 9-TB-Dox treatment (**Fig. 1C & 1D**). NCAPD3 immunostaining of the same tissue section revealed broad NCAPD3 expression in all cells of control mice and decreased NCAPD3 staining intensity in the GFP-positive epithelium of 9-TB-Dox-treated mice. GFP expression varied along the crypt axis with lower/no expression in the crypt base and strong expression closer to the crypt tip, mirroring reported increased expression of villin as epithelial cells differentiate (21). Importantly, the GFP and NCAPD3 staining did not overlap, indicating effective knockdown in shRNA-expressing cells. To more directly assess the degree of NCAPD3 knockdown in intestinal epithelial cells after transgene induction, flow-sorted colonic EpCAM^+^/ CD45^-^/ GFP^+^ cells from 9-TB-Dox mice were compared to EpCAM^+^/ CD45^-^ cells from control mice. Immunoblot analysis of NCAPD3 protein levels revealed a marked reduction (75%) in intestinal epithelial cells from 9-TB-Dox treated mice (**Fig. 1E**). These data demonstrate the successful genetic engineering of mice to create an inducible, intestinal epithelial cell-specific knockdown of NCAPD3 expression to facilitate the *in vivo* evaluation of the role of NCAPD3 in the intestinal epithelium.

### Short term NCAPD3 knockdown in the intestinal epithelium is not sufficient to induce spontaneous colitis

To assess the direct impact of reduced NCAPD3 expression in intestinal epithelial cells *in vivo*, NCAPD3KD mice were treated daily with 9-TB-Dox or saline for 10 days. Although 9-TB-Dox-treated mice tended to lose more weight than control animals, no significant differences in weight loss, gross colonic appearance, or colon length were observed (**Fig. 2A & 2B**). Colonic tissue histopathology evaluation using a standardized colitis scoring system did not identify signs of intestinal inflammation in any animal (**Fig. 2C**). Intestinal permeability was assessed using a flagellin bioassay on endpoint plasma samples and showed a trend of increased permeability in NCAPD3KD mice treated with 9-TB-Dox (**Fig. 2D**). In addition, 9-TB-Dox treated NCAPD3KD mice had 2.1 fold increased levels of the systemic acute phase marker of inflammation, SAA relative to control mice (**Fig. 2E**). Overall, these data indicate that short term NCAPD3 knockdown in the intestinal epithelium is not sufficient to induce spontaneous colitis but appears to mildly elevate basal inflammation and gut permeability.

**Figure 2:**
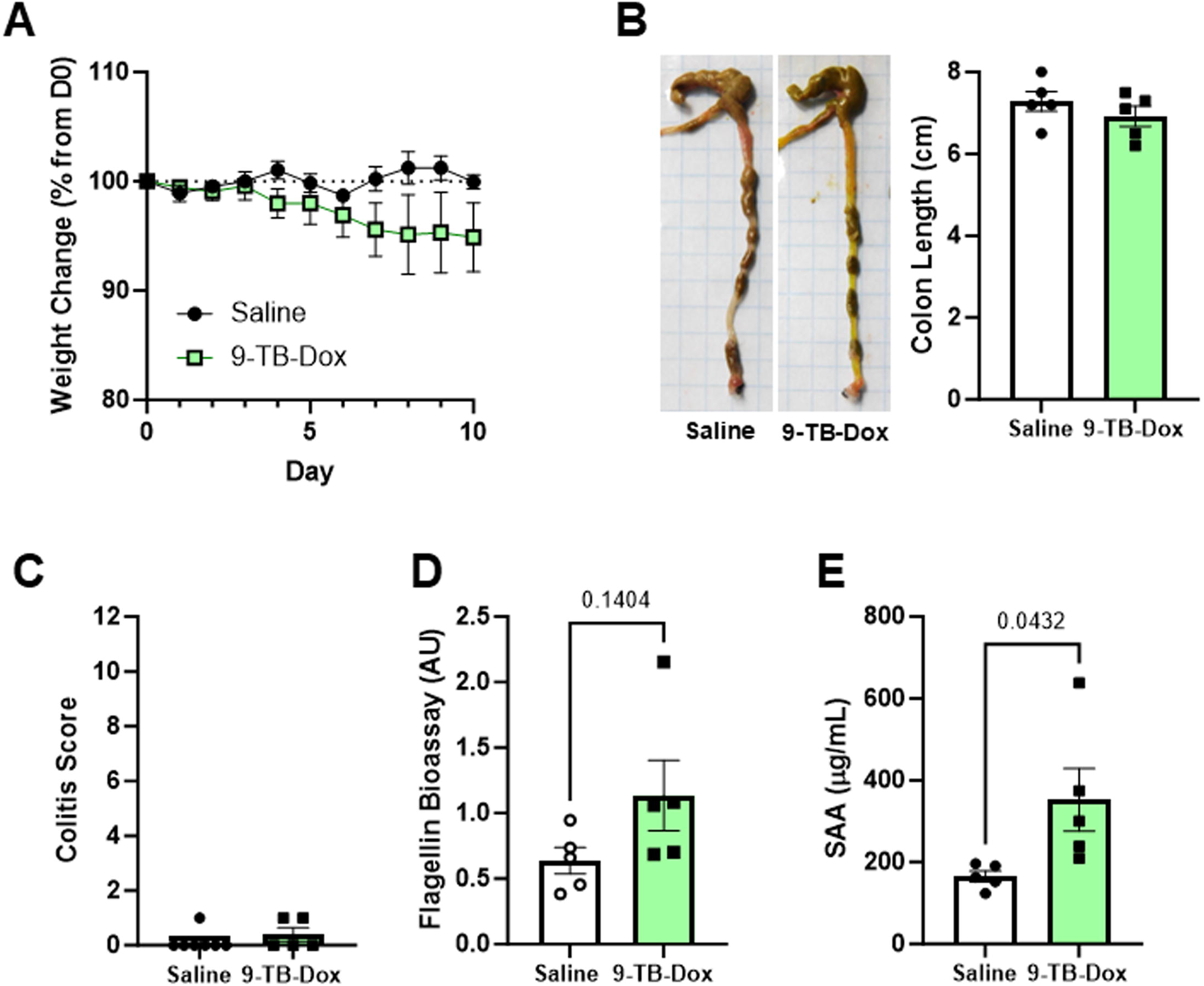
Intestinal epithelial NCAPD3 knockdown does not result in spontaneous colitis. Mice were treated with saline or 9-TB-Dox (25 mg/kg i.p.) daily for 10 days (n=5 mice/group). **A**. Weight change relative to starting weight. **B**. Gross evaluation of colonic pathology at endpoint. **C**. Histopathology of distal colon tissue. **D**. Intestinal barrier integrity assessed by plasma bioassay on HEKBlue-mTLR5 reporter cell line. **E**. Systemic inflammation assessed by quantification of SAA at endpoint. Mean ±SEM graphed. Significance determined by unpaired 2-tailed t-test.

### Epithelial NCAPD3 knockdown increases sensitivity to intestinal damage induced by DSS

To further investigate the role of NCAPD3 in intestinal barrier maintenance, NCAPD3KD mice were treated with or without 9-TB-Dox and administered low-dose DSS for 7 days to induce mild intestinal injury. NCAPD3KD mice treated with 9-TB-Dox lost more weight (**Fig. 3A**) and had a higher DAI (**Fig. 3B & Table S2**) than saline-treated controls. Quantification of colonic tissue histopathology revealed increased colitis in 9-TB-Dox-treated mice (**Fig. 3C & Table S3**), driven primarily by increased epithelial damage scores (2.17 saline vs. 2.80 9-TB-Dox; p=0.02). Correlating with increased inflammation and intestinal epithelium damage were increased levels of the proinflammatory chemokines, MIP1α/CCL3 and MIP2/CXCL2, as well as pro-inflammatory, epithelial damaging cytokines, IL-9, IL-15, and IL-17, in colonic tissue samples (**Fig. S3**). Intestinal permeability of 9-TB-Dox treated mice was approximately double that of control mice, as measured by 4 kDa FITC-dextran transit to the blood after oral challenge (**Fig. 3D**) and by flagellin bioassay of endpoint plasma samples (**Fig. 3E**). Endpoint SAA levels were also dramatically elevated 3.8-fold in 9-TB-Dox mice relative to controls (**Fig. 3F**). These results indicate that reduced NCAPD3 expression in intestinal epithelial cells sensitizes mice to intestinal injury by the chemical irritant DSS.

**Figure 3:**
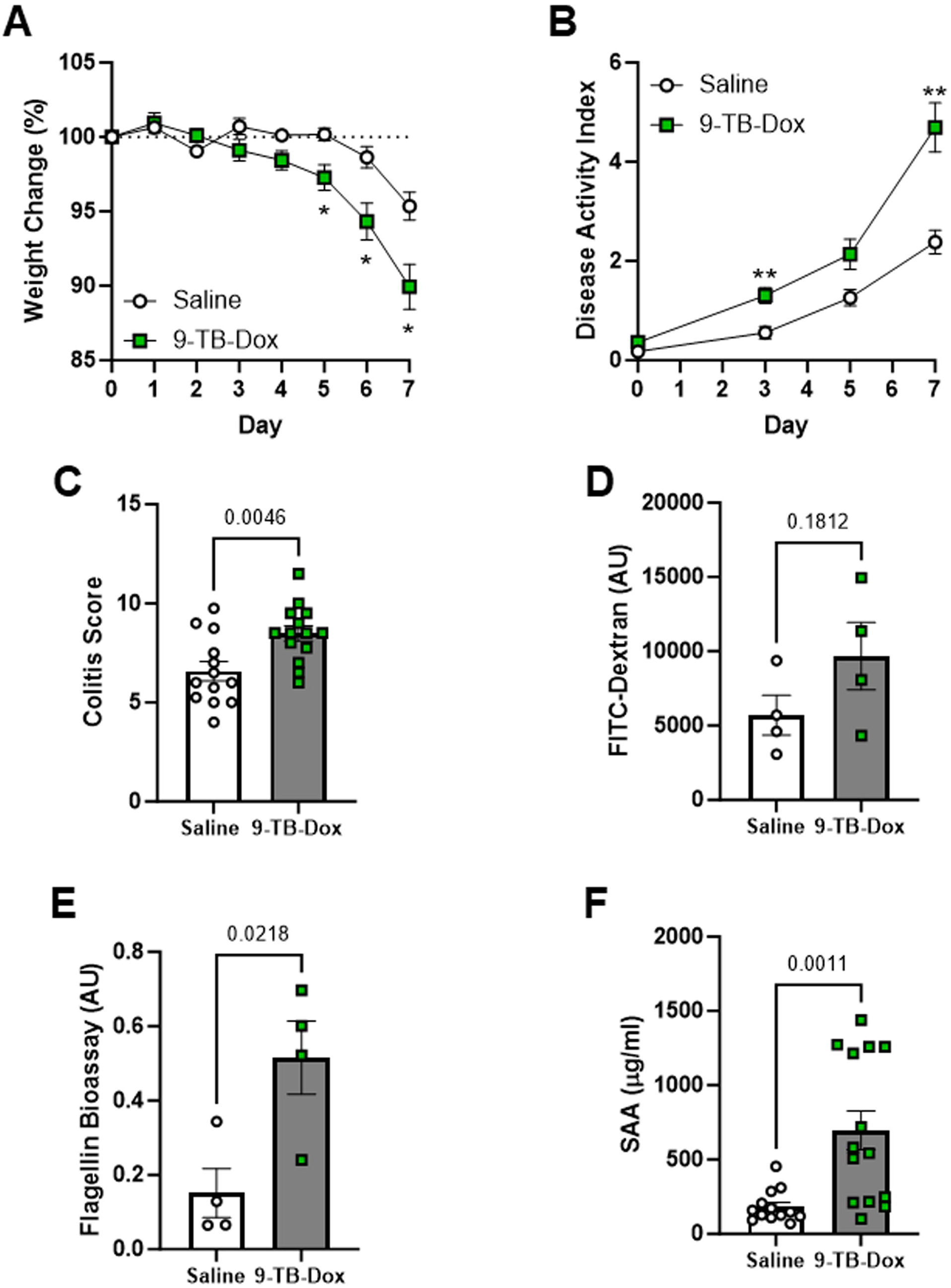
Knockdown of NCAPD3 expression in intestinal epithelial cells sensitizes mice to intestinal injury with dextran sodium sulfate. NCAPD3 KD mice (n=16-18 mice/group) were treated daily with either saline or 9-TB-Dox (25 mg/kg i.p.) 4 days prior to 1.5% DSS administration for 7 days. **A**. Weight change during DSS treatment. **B**. Disease activity index during DSS treatment. **C**. Evaluation of distal colon histopathology using a validated colitis scoring system. **D**. Paracellular intestinal permeability monitored by serum fluorescence levels 2 hours after 4 kDa FITC-dextran gavage (12 mg/mouse; n=4 mice/group). **E**. Measurement of intestinal permeability by flagellin bioassay using endpoint serum samples (n=4 mice/group). **F**. Systemic inflammation evaluated by SAA levels. Mean± SEM graphed. Significance determined by 2-way ANOVA with Sidak’s multiple comparisons test (A. & B.) or 2-tailed unpaired t-test (C.-F.).

### NCAPD3 expression strengthens epithelial defense against the diarrheal pathogen *Salmonella*

Our prior studies in human intestinal epithelial cell culture and *Drosophila* model systems demonstrated a role for NCAPD3 in the upregulation of antimicrobial defenses in response to bacterial pathogen challenge (10, 11). Therefore, we examined the impact of NCAPD3 knockdown on the resistance of mice to oral challenge with the diarrheal pathogen *Salmonella enterica serovar* Typhimurium. As infection of C57BL/6 mice with wild-type *S*. Typhimurium results in acute systemic typhoid disease, we used the attenuated ΔAroA strain to cause a chronic enteric infection to better replicate human diarrheal disease (22).

First, we characterized the impact of 9-TB-Dox on the growth and survival of *Salmonella* ΔAroA. Using an *in vitro* growth assay, we compared the bactericidal activity of 9-TB-Dox to doxycycline (**Fig. S4A**). After 20 hours, doxycycline induced a dose-dependent decrease in *Salmonella* growth, while 9-TB-Dox had no effect. In contrast, a pilot mouse *Salmonella* ΔAroA infection of WT C57BL/6 mice treated with or without 9-TB-Dox showed no differences in fecal bacterial shedding, but increased colonization of cecum and colon tissues (**Fig. S4B-D)**. Therefore, we modified our original experimental design to treat all mice with 9-TB-Dox and compare NCAPD3KD mice (KD) with founder mice lacking the rtTA driver required for induction of the transgene cassette (control; C) in the *Salmonella* ΔAroA oral infection model.

All mice were pre-treated with 9-TB-Dox, then orally challenged with *Salmonella* ΔAroA, and monitored for 7 days post-infection. Weight loss post-infection and fecal *Salmonella* shedding were moderately higher in KD mice (**Fig. 4A & 4B**). However, no differences were observed in the tissue levels of viable *Salmonella* in the cecum, colon, mesenteric lymph nodes, liver, or spleen (**Fig. 4B**). Epithelial knockdown of NCAPD3 resulted in increased gross colonic damage quantified by a diarrheal scoring system (**Fig. 4C & Table S4**). Further examination of intestinal tissues revealed increased histopathology scores in both the cecum and colon of KD mice (**Fig. 4D & Table S5**). The increased tissue damage correlated with diminished expression of the antimicrobial peptides Reg3β and defensin-β14 (DEFB14) in KD mice (**Fig. 4E**). Together, this shows a role for NCAPD3 in maintenance of intestinal homeostasis during enteric *Salmonella* infection.

**Figure 4:**
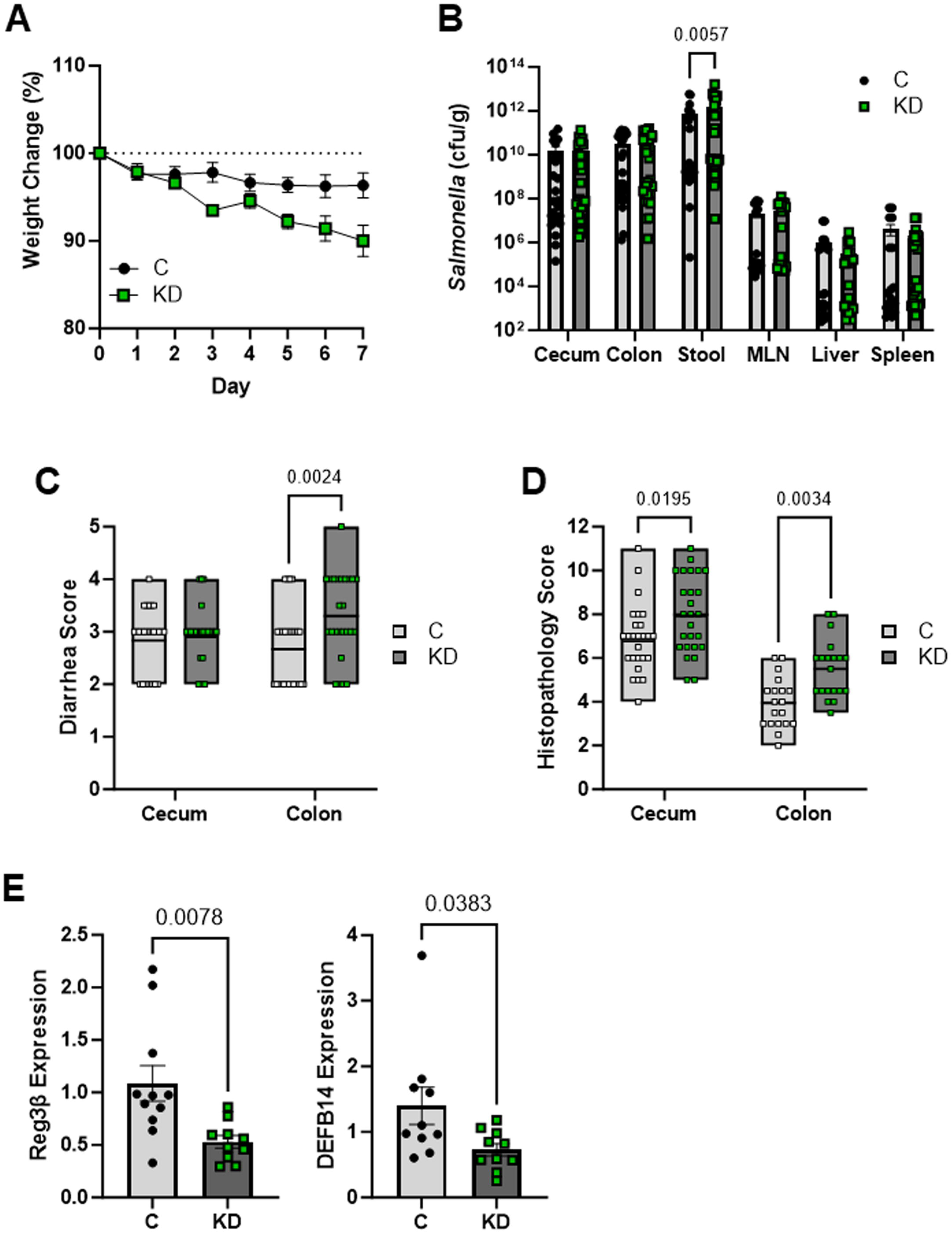
NCAPD3 KD mice are more susceptible to intestinal damage from *Salmonella* Typhimurium infection. NCAPD3 KD mice (KD) and mice that lacked the villin-rtTA transgene (C) were treated daily with 9-TB-Dox (25 mg/kg) for 4 days prior to oral antibiotic treatment and subsequent oral infection with *Salmonella enterica* serovar Typhimurium ΔAroA (n=25-28 mice/group). **A**. Weight change after *Salmonella* infection (p=0.0145). **B**. Tissue and stool bacterial loads. **C**. Endpoint gross diarrhea score. **D**. Endpoint histopathology score. **E**. Cecal transcript levels of antimicrobial peptides. Transcripts were normalized to 18S and graphed relative to the mean of control (C) mice. Mean ±SEM graphed. Significance determined by 2-way ANOVA with Sidak’s multiple comparisons test (A.-D.) or unpaired 2-tailed t-test (E.).

## Discussion

In this study, we generated and validated a novel inducible, intestinal epithelial cell-specific NCAPD3 knockdown mouse model and used it to investigate the *in vivo* role of NCAPD3 in intestinal barrier homeostasis. Using a villin-rtTA driven shRNA system targeted to the Col1a1 safe harbor locus, we achieved robust epithelial-specific reduction of NCAPD3 following induction with the non-antimicrobial tetracycline analogue 9-TB-Dox. This *in vivo* mammalian model enables mechanistic investigation of the observation that UC patients have reduced NCAPD3 expression specifically in intestinal epithelial cells (11). Our data support the interpretation that this epithelial NCAPD3 reduction is not a secondary consequence of inflammation but may contribute to epithelial dysfunction and disease pathogenesis.

Despite efficient (∼75%) epithelial knockdown over 10 days, NCAPD3KD mice did not develop spontaneous colitis. However, the trend toward increased intestinal permeability, elevated circulating flagellin levels, and significantly increased SAA suggest that NCAPD3 loss subtly compromises epithelial barrier integrity, even in the absence of overt histological inflammation. Increased intestinal permeability is increasingly recognized as an early event in IBD, detectable even in quiescent disease and in individuals prior to clinical onset (23-25). The observation that NCAPD3KD mice exhibit low-level permeability and systemic inflammation without visible tissue injury supports the concept that NCAPD3 loss can contribute to epithelial barrier weakening and predispose one to inflammatory conditions. While this study examined only short-term knockdown, chronic NCAPD3 deficiency may exacerbate this basal inflammatory tone and potentially lead to spontaneous disease. Extended knockdown studies will be required to determine whether long-term NCAPD3 reduction alone is sufficient to break the intestinal barrier and drive intestinal pathology.

The increase in SAA in NCAPD3KD mice, even at baseline, is notable. SAA is not merely a marker of inflammation but an active participant in inflammatory processes, capable of inducing cytokine production, recruiting immune cells, and contributing to tissue repair responses (26). In this context, elevated SAA in the absence of overt injury may reflect ongoing low-level epithelial stress and induction of repair mechanisms triggered by a subtle compromise to the barrier. Importantly, SAA levels were further amplified following DSS exposure, consistent with an exaggerated systemic response when the compromised epithelium encounters an additional insult.

When challenged with DSS or an enteric *Salmonella* infection, NCAPD3KD mice exhibited significantly heightened disease activity, epithelial damage, inflammatory cytokine production, and intestinal permeability compared to controls. The amplification of permeability and inflammation in these settings aligns with the widely proposed “two-hit hypothesis” for IBD pathogenesis, where a genetically predisposed individual may have epithelial vulnerability and mild barrier dysfunction that, when coupled with environmental triggers, can lead to an uncontrolled inflammatory state (27). In this context, reduced NCAPD3 expression may represent a susceptibility factor that does not independently cause disease but sensitizes the host to secondary insults.

During DSS-induced injury, NCAPD3 deficiency was associated with significantly increased epithelial damage and elevation in the proinflammatory mediators MIP1α/CCL3, MIP2/CXCL2, IL-9, IL-15, and IL-17. IL-17 and IL-15 are well-established contributors to mucosal inflammation in both DSS colitis and human IBD, promoting leukocyte recruitment and sustaining immune activation (28). Notably, MIP1α/CCL3 and MIP2/CXCL2 are key chemokines in IBD that promote the recruitment of macrophages and neutrophils, respectively, and their upregulation correlates with disease severity (29). Whilst it remains unclear whether these changes reflect direct transcriptional dysregulation resulting from NCAPD3 loss or are secondary to increased epithelial damage and permeability, this data suggests that NCAPD3 plays a role in restraining inflammatory amplification following barrier disruption. Further studies are required to determine whether this occurs through direct chromatin-mediated regulation of inflammatory gene programs or indirectly because of worsened epithelial injury.

Consistent with our prior *in vitro* and *Drosophila* studies, epithelial NCAPD3 knockdown compromised host defense during *Salmonella* infection. Although tissue bacterial burdens were comparable, NCAPD3KD mice exhibited increased diarrheal severity and worsened histopathology. This was accompanied by reduced expression of the antimicrobial peptides Reg3β and DEFB14. Reg3β is strongly induced during a *Salmonella* enteric infection and contributes to maintaining a distance between luminal bacteria and the epithelial surface (30), while β-defensins directly limit bacterial growth and shape microbiota composition (31). Deficits in antimicrobial peptide production also have implications for IBD, as antimicrobial peptide levels can be reduced even in the presence of pro-inflammatory signaling that typically upregulates their expression (32). As prior enteric infections are known to increase IBD risk in genetically predisposed individuals (33), heightened tissue damage during infection in the setting of reduced NCAPD3 may represent another mechanism linking epithelial chromatin dysregulation to chronic inflammatory susceptibility.

Recent advances have highlighted broader roles for chromatin regulators in controlling transcriptional programs governing epithelial differentiation and innate immunity (34). For example, the chromatin architectural protein CTCF, together with cohesin, organizes higher-order chromatin looping at inflammatory cytokine loci and directly regulates transcriptional responses (35, 36). Similarly, SATB1 acts as a genome organizer in immune cells, controlling the expression of cytokines and other immune genes by regulating long-range chromatin interactions (37, 38). NCAPD3 also contributes to genomic stability by repressing retrotransposons and limiting apoptosis (8, 39). Loss of this function can increase DNA damage and cell stress, potentially weakening epithelial turnover and barrier integrity. These examples demonstrate how disruption of chromatin-organizing systems can reprogram inflammatory, stress-response, and differentiation pathways, indicating mechanisms by which dampened levels of NCAPD3 could reshape epithelial function through skewing expression of cytokines, antimicrobial peptides, or junctional proteins.

In summary, our study establishes NCAPD3 as an epithelial regulator of the mammalian intestine that enhances epithelial barrier resilience and antimicrobial defense during stress. Although dispensable for short-term basal homeostasis, NCAPD3 function becomes critical during epithelial injury and enteric infection. Reduced NCAPD3 expression may therefore lower the threshold for inflammatory disease by weakening barrier integrity, amplifying inflammatory cascades, and impairing antimicrobial defenses. These findings position NCAPD3 as a potential modulator of IBD susceptibility and highlight chromatin organization as an important, previously underappreciated layer of intestinal epithelial regulation.

## Materials and Methods

### NCAPD3 KD transgenic mouse line generation

All animal experiments were conducted in accordance with institutional and national guidelines. Animal use protocols were reviewed and approved by the Institutional Animal Care and Use Committees (IACUC) of Cleveland Clinic and Case Western Reserve University. The genetic manipulation of mice was reviewed and approved by the Institutional Biosafety Committee (IBC) of Case Western Reserve University. Experiments were conducted following the ARRIVE guidelines.

Short hairpin RNA (shRNA) sequences targeting mouse NCAPD3 were tested for knockdown of NCAPD3 protein by generating stable immortalized B6 mouse macrophage cell lines (40) transduced with isopropyl β-D-1-thiogalactopyranoside (IPTG)-inducible shRNA lentiviral constructs (pLKO-puro-IPTG-3xLacO; Sigma; **Supplemental Table 1**). Expression of the shRNA was induced by adding 2 mM IPTG to the cell culture medium for 1-3 days. NCAPD3 protein levels relative to a loading control (Na/K ATPase) were determined by densitometric analysis of cell lysate immunoblots using ImageJ software (41). Degree of knockdown was determined by comparison to cells expressing a non-targeting shRNA treated with IPTG (**Fig. S1A**).

CRISPR/Cas targeting was used to insert a NCAPD3-targeting shRNA and green fluorescent protein (GFP) expression cassette into the safe harbor locus of Col1a1 (42). A guide sequence, Col1a1 607/rev (CAGGCCTCAGGTCTGCAGCA), that directs cutting near the PstI site downstream of the Col1a1 3’-UTR, was validated using the Guide-it Complete sgRNA Screening System (Clontech) against a genomic target amplified from B6SJL/F1 DNA. Large-scale preparations of Col1a1 607/rev sgRNA and Cas9 nuclease were ordered from PNABio and prepared as directed.

A targeting vector, incorporating 1 kb 5’ and 3’ arms of homology from Col1a1, was designed to insert the NCAPD3-targeting shRNA expression cassette at the position of the double-strand break directed by the guide. The NCAPD3-targeting shRNA cassette was closely modeled on the cTGM vector from the Scott Lowe laboratory (43), but without the elements required for embryonic stem cell selection. The vector was designed using VectorBuilder.com and ordered as a large-scale supercoiled preparation (VectorBuilder). Aliquots of the vector were spin clarified twice through Ultrafree-MC filters (Millipore).

Mixtures of Cas9 nuclease, sgRNA, and vector were prepared in Nuclease Free water (Qiagen) and injected into the pro-nuclei of 0.5 dpc B6SJL/F2 fertilized oocytes. The mixtures, in ng/µl of Cas9/sgRNA/vector, ranged from 20/20/2 to 5/5/0.5. Injected embryos were allowed to recover and surgically transferred through the oviduct into 0.5 dpc CD1 pseudo-pregnant recipients. The resulting 78 pups were transferred at weaning from the Case Transgenic and Targeting facility (Case Western Reserve University) to the principal investigator’s laboratory at the Cleveland Clinic.

The F_0_ founders (both males and females) were initially screened by PCR for the presence of the NCAPD3-targeting shRNA expression cassette using primers internal to the targeting vector. The correct targeting of cassette-positive animals was then verified by LR-PCR using external and internal primers on both sides and verified by sequencing of the LR-PCR products. Selected correctly targeted males and females were back crossed to wild-type mice to generate three founder lines (**Fig. S1B**). These lines were crossed to transgenic mice with villin-1 regulatory elements driving expression of an optimized reverse tetracycline-controlled transactivator protein (rtTA*M2) (Jackson Labs strain#031285; B6;SJL-Tg(Vil1-rt-TA*M2)5Stang/J) to result in a strain with tetracycline/doxycycline-inducible expression of the NCAPD3-targeting shRNA cassette in the intestinal epithelium. These double-transgenic lines (NCAPD3 KD mice) were backcrossed 5-7 generations onto a C57BL/6J strain background (Jackson Labs strain#000664), alternating male and female backcross partners. Carriage of NCAPD3-targeting shRNA expression cassette and villin-rtTA*M2 transgene determined by a real-time TaqMan PCR using custom 3 probe panel (Transnetyx Inc.). As the phenotypes of the three NCAPD3 KD lines were similar, results from all three lines were combined in the data presented.

NCAPD3 knockdown was induced through daily intraperitoneal (i.p.) injection of 25 mg/kg 9-tert-butyl doxycycline HCl (9-TB-dox; Echelon Biosciences, cat# B-0801). Expression of the transgene cassette was monitored routinely by detection of GFP-expressing tissues using a handheld UV light source or IVIS Spectrum CT small animal imager (PerkinElmer). NCAPD3 protein knockdown was confirmed by immunofluorescent confocal microscopy and immunoblot of flow-sorted intestinal epithelial cells.

### Immunofluorescent staining of intestinal tissues

Slides with tissue sections 8-10 µm thick were prepared from fresh frozen tissue embedded in OCT. After thawing at room temperature for 15-30 minutes, slides were incubated in ice cold methanol for 10 minutes, followed by three washes in phosphate buffered saline (PBS) for 5 minutes each on a shaker. Slides were incubated in blocking buffer (5% human serum albumin/PBS) in a humid chamber at room temperature for 30-60 minutes. Tissue sections were then incubated with NCAPD3 antibody (Bioss, cat# bs-7734R, 1:100 dilution) in blocking buffer overnight at 4°C. Slides were washed three times in PBS, and goat anti-mouse IgG conjugated to Alexa Fluor 488 (ThermoFisher, cat# A11001) diluted in blocking buffer was added. Slides were placed in a dark humid chamber for one hour and then washed three times with PBS. Coverslips were mounted with Vectashield with DAPI (Vector Labs) and sealed with nail polish. Images were obtained using a Leica TCS-SP8-AOBS confocal microscope equipped with LAS-X software (Leica Microsystems).

### Flow cytometry enrichment of intestinal epithelial cells

Mice were treated daily for 4 days with 25 mg/kg 9-TB-dox, (i.p.). Colons were removed, filleted open, and washed three times with ice-cold phosphate-buffered saline (PBS). Tissue was incubated on ice for 20 min in 30 mM EDTA/ 1.5 mM DTT. Tissue was then transferred to 30 mM EDTA and incubated at 37°C for 10 min. Epithelial cells were dislodged from bulk tissue by shaking, as previously described (44, 45). Cells were pelleted by centrifugation at 100 x g for 3 min at 4°C, washed with ice-cold PBS, and pelleted again. Cells were incubated in PBS with 8 mg/ml dispase (Gibco cat# 17105-041) and 5 mg/ml DNase (Worthington, cat# LS002007) at 37°C for 20 min. Additional PBS was added, cells were passed through a 70 µm cell strainer, and then pelleted at 100 x g at 4°C for 3 min. Cells were resuspended in LIVE/DEAD Fixable Violet Dead Cell Stain (ThermoFisher cat# L34964) and incubated at 4°C for 20 min. Cells were washed with ice-cold PBS and centrifuged at 100 x g at 4°C for 3 min. Stained cells were incubated with FcR blocking reagent (Miltenyi Biotec cat#130-092-575) for 5 min at 4°C and then subsequently stained with APC-conjugated anti-CD45 (1:200, Biolegend, Cat#109814) and PE-conjugated anti-EpCAM/CD326 (1:50, ThermoFisher cat# 12-5791-83) at 4°C for 30 min. Cells were washed with PBS/ 1 mM EDTA/ 0.5% BSA and pelleted at 100 x g at 4°C for 3 min. Cells were resuspended in PBS/ 2 mM EDTA/ 2% FBS, passed through a 35 µM filter, and sorted on a Bigfoot Spectral Cell Sorter (ThermoFisher) for the collection of viable CD45^-^ /Epcam^+^ epithelial cells, further categorized by GFP expression. Gating strategy shown in **Fig. S2**.

### Dextran sodium sulfate colitis model

For colitis experiments, 7-11 week old male and female mice were randomized into treatment groups with littermates distributed between treatment groups whenever possible. Mice were i.p. injected with 9-TB-dox (25 mg/kg) daily beginning 4 days prior to colitis induction and throughout the experiment. Colitis was induced over 7 days through the addition of 1.5% dextran sulfate sodium (DSS; 36,000–50,000 m.w., MP Biomedicals, LLC) to the drinking water. Mice were monitored daily to calculate a disease activity index score (DAI; **Table S2**) and sacrificed on day 7 for harvest of blood and tissues. Serum amyloid A (SAA) levels were measured from plasma by ELISA (Immunology Consultants Laboratory). Portions of terminal ileum and proximal, transverse and distal colon were fixed in 10% neutral buffered formalin for histologic analyses. Hematoxylin and eosin-stained sections were evaluated blinded to treatment group and genotype to determine a colitis score (**Table S3**). Distal colon segments were collected for analysis of cytokines/chemokines.

### Salmonella colitis model

For colitis experiments, 7-11 week old male and female mice were randomized into treatment groups based on genotype with littermates distributed between treatment groups whenever possible. Mice were i.p. injected with 9-TB-dox (25 mg/kg) every 24 hours beginning 4 days prior to infection and throughout the experiment. Twenty-four hours prior to infection, mice were orally gavaged with 20 mg streptomycin. *Salmonella enterica* Typhimurium ΔAroA SL3261 (gift from Bobby Cherayil, Department of Pediatrics, Harvard Medical School, Boston, MA) was grown at 37°C with shaking in LB Miller broth supplemented with 50 µg/ml streptomycin. For infections, a stationary culture diluted in PBS was used. Food was withdrawn for 4 hours prior to oral gavage with 3×10^7^ cfu *Salmonella enterica* Typhimurium ΔAroA SL3261 in 100 µl of PBS. Mice were weighed and monitored daily and sacrificed on day 7 for harvest of blood and tissues. A gross evaluation of diarrhea and tissue damage was taken during dissection using a modified diarrhea score (46) (**Table S4**). SAA levels were measured from plasma by ELISA (Immunology Consultants Laboratory). For bacterial enumeration, tissues (liver, spleen, mesenteric lymph nodes (MLN), cecum, and colon) and stool were collected in 1 ml of sterile PBS, stored on ice, and homogenized with an Omni Tissue Homogenizer (Omni International). Homogenized tissues were serially diluted and plated onto LB agar plates containing 50 µg/ml streptomycin and incubated overnight at 37°C. Calculated CFUs were then normalized to the weight of the tissue to obtain CFU/g. Portions of the terminal ileum, cecum, and proximal and distal colon were fixed in 10% neutral buffered formalin for histological analyses. Hematoxylin and eosin-stained sections were evaluated to determine the histopathology score (47) (**Table S5**).

### Intestinal permeability assays

For the determination of intestinal permeability by small molecule tracer flux *in vivo*, mice were food-restricted for 3 hours prior to oral gavage with 12 mg fluorescein isothiocyanate-dextran (FITC; average MW: 3000-5000, Sigma Aldrich) suspended in 100 µL sterile water. Blood was collected 2 hours post-gavage, and serum was separated by centrifugation for 5 min at 5,000 rpm. Plasma was diluted in PBS and fluorescence determined at 530 nm with excitation at 485 nm. Measurement of intestinal permeability by flagellin bioassay was performed by adding diluted plasma in duplicate to HEKBlue-mTLR5 reporter cells (InVivoGen) plated at 2.5 x10^4^ cells/well in 96-well plates for 24 hours. Cell culture supernatants were assayed for secreted alkaline phosphatase using QUANT-Blue (InVivoGen) solution at 37°C for 3 hours.

### Quantification of cytokines/chemokines from intestinal tissue

Colon and cecal tissues were manually homogenized in lysis buffer (20 mM Tris pH 7.5, 150 mM NaCl, 1mM EDTA, 1mM EGTA, 1% Triton X100) with phosphatase and protease inhibitors (Sigma). Protein in the tissue homogenates was quantified using a BCA assay (Pierce) and normalized to 25 μg/well. Multiplexed cytokine and chemokine analysis was performed using the Proinflammatory Panel 1 (mouse) and Cytokine Panel 1 (mouse) kits from Meso Scale Diagnostics (MSD; Cat #K15048D, #K15245D) following the manufacturer’s instructions. The panels included the following analytes: IFN-γ, IL-1β, IL-2, IL-4, IL-5, IL-6, KC/GRO, IL-10, IL-12p70, TNF-α, IL-9, MCP-1, IL-33, IL-27p28/IL-30, IL-15, IL-17A/F, MIP-1α, IP-10, MIP-2. Data was analyzed using the MSD software, MSD Discovery Workbench 4.0.

### Immunoblots

For analysis of NCAPD3 protein expression level, cells were lysed in Laemmli buffer (310 mM Tris pH 6.8, 25% glycerol, 5% SDS, 715 mM 2-mercaptoethanol, 125 mg/mL bromophenol blue) on ice. The resulting lysate was separated by SDS-PAGE and proteins transferred to a PVDF membrane using an iBlot2 transfer system (Invitrogen). Membranes were blocked in 5% milk/Tris-buffered saline with 0.1% Tween-20 (TBST) and then probed with primary antibodies (anti-CAPD3, Bethyl, A300-604A 1:1000, or anti-α-Tubulin, Sigma, T9026, 1:20,000) overnight at 4°C. After washing with TBST, blots were probed with HRP-conjugated secondary antibodies (Jackson Laboratories) for 1 hour in 2.5% milk/TBST. Blots were developed by enhanced chemiluminescence (Millipore). Uncropped blots used in figures shown in **Fig. S5**.

### Antibiotic testing of 9-TB-Dox

To determine the antimicrobial effects of 9-TB-dox, log phase *Salmonella enterica* serovar Typhimurium ΔAroA SL3261 cultures were diluted into LB medium with the indicated concentrations of 9-TB-dox or doxycycline in 96-well plates in triplicate. Bacterial growth over time at 37°C was monitored by absorbance at 600 nm in a BioTek Synergy H1 plate reader.

### Quantitative Real Time PCR (qRT-PCR)

Total RNA was isolated using Trizol (ThermoFisher) from total cecal tissue homogenates, per manufacturer’s instructions. RNA was converted to cDNA using iScript cDNA synthesis kit (BioRad), containing 1 µg RNA per reaction. Transcript levels were measured by qPCR (see **Table S6** for primer information) in duplicate using SYBR Green supermix (BioRad) and quantified using the 2^-ΔΔCT^ method.

## Statistical Analysis

Statistical analyses were performed using GraphPad Prism software (version 10). Groups were compared by unpaired 2-tailed t-test or one-way ANOVA with *post-hoc* Sidak’s multiple comparisons test. Differences were considered significant when p<0.05. Specific tests used for each dataset are detailed in the figure legends.

## Data Availability

All relevant data are publicly available within the manuscript and its supporting information files.

## Acknowledgments

We thank David LePage and Ron Conlon in the Case Transgenic & Targeting Facility for their guidance, design, and generation of the targeted NCAPD3KD mouse founders. We appreciate the assistance of Dr. Kewal Asosingh and the expertise of Cleveland Clinic Research Flow Cytometry Core Facility (RRID:SCR_026460) in optimization of the isolation and analysis of intestinal epithelial cells. We also thank Apryl Helmick in the Cleveland Clinic Research Histology Core for patiently processing all our mouse tissues. We are very appreciative of the technical support and guidance from members of the McDonald laboratory, Andràs Ponti, Lily Markley, and Isabella Willis, as well as Dr. Kim Such and Tori Evans from the Biological Resources Unit. We thank Dr. Bobby Cherayil for the gift of the *Salmonella* ΔAroA strain. We especially thank Dr. Serpil Erzurum for her timely support of this project and ongoing support of our research programs.

