## Supplementary Materials File for "Epithelial NCAPD3 expression protects against stress-induced intestinal injury in mice"

#### Supplemental Tables

**Table S1: Sigma MISSION shRNA constructs**

| TRC Clone ID | Target Region | NCAPD3 KD[1] | Sequence |
|---|---|---|---|
| TRCN0000199006 | Coding | 10% | CCGGGCTCTCAGATACCTTTGACATCTCGAGATGTCAAAGGTATCTGAGAGCTTTTTTG |
| TRC0000432911 | 3'UTR | 0% | CCGGTATACTGTGATACTCTATTTGCTCGAGCAAATAGAGTATCACAGTATATTTTTTG |
| TRC0000215444 | 3'UTR | 60% | CCGGGACTTTCTCACAATAACATTACTCGAGTAATGTTATTGTGAGAAAGTCTTTTTTG |
| SHC312V | Non-target | 0% | CCGGGCGCGATAGCGCTAATAATTTCTCGAGAAATTATTAGCGCTATCGCGCTTTTT |

[1]Percent decrease of NCAPD3 protein levels compared to non-target control cells after 48 hours IPTG induction

**Table S2: Dextran Sulfate Sodium Colitis Disease Activity Score**

| Component | Score | Definition |
|---|---|---|
| Weight Loss | 0 | 0 – 5% |
| | 1 | >5 – 10% |
| | 2 | >10 – 15% |
| | 3 | >15 – 20% |
| Posture | 0 | Normal |
| | 1 | Hunched |
| Fur Appearance | 0 | Smooth |
| | 1 | Ruffled |
| Stool | 0 | Normal |
| | 1 | Soft |
| | 2 | Soft with Blood |
| | 3 | Runny with Blood |
| Rectal Prolapse | 0 | None |
| | 1 | 1 mm |
| | 2 | 2 mm |

**Table S3: Dextran Sulfate Sodium Colitis Score**

| Component | Score | Definition |
|---|---|---|
| Epithelium | 0 | intact epithelial layer, well defined crypt structure |
| | 1 | intact epithelial layer, reduced crypt structure |
| | 2 | some breaks in epithelium, reduced/loss of crypt structure |
| | 3 | loss of epithelium & loss of crypt structure |
| Leukocyte Infiltration | 0 | none |
| | 1 | sparse |
| | 2 | moderate |
| | 3 | fulminant |
| Submucosal Swelling | 0 | none |
| | 1 | minor |
| | 2 | moderate |
| | 3 | severe |
| Muscularis Hyperplasia | 0 | none |
| | 1 | minor |
| | 2 | moderate |
| | 3 | severe |

**Table S4: *Salmonella* Gross Diarrhea Score[1]**

| Tissue | Score | Definition |
|---|---|---|
| Colon | 0 | normal colon with formed feces in the distal colon or rectum |
| | 1 | areas of white tissue and/or gas bubbles in the lumen; the rest of the colon is normal |
| | 2 | involvement of the proximal colon (e.g. fluid filled); formed stools in distal colon |
| | 3 | score of 2 plus fluid throughout most of the colon; soft feces in the distal colon |
| | 4 | no formed feces in the entire colon; colon filled with clear or bloody mucus |
| Cecum | 0 | normal cecum |
| | 1 | cecum that is smaller than normal with areas of white tissue and stool contents |
| | 2 | severe typhlitis (white, shrunken cecum) very minimal stool |
| | 3 | severe typhlitis (white, shrunken cecum) no contents |
| | 4 | severe typhlitis (white, shrunken cecum), contains bloody mucus |

[1]modified from Woo et al., 2008 (1)

**Table S5: *Salmonella* Histopathology Score[1]**

| Component | Score | Definition |
|---|---|---|
| Submucosal Swelling | 0 | none |
| | 1 | minor <0.2mm wide <50% diameter |
| | 2 | moderate 0.2-0.45mm, 50-80% |
| | 3 | severe >0.46mm, >80% |
| PMN Infiltration[2] | 0 | <5 |
| | 1 | 5-20 |
| | 2 | 21-60 |
| | 3 | 61-100 |
| | 4 | >100 |
| Goblet Cells[3] | 0 | >28 |
| | 1 | 11-28 |
| | 2 | 1-10 |
| | 3 | <1 |
| Epithelium | 0 | intact epithelial layer, well defined crypt structure |
| | 1 | epithelial desquamation |
| | 2 | epithelial erosion (gaps of 1-10 cells) |
| | 3 | epithelial ulceration (gaps of >10 cells) |

[1]modified from Barthel et al., 2003 (2)
[2]average number of PMN observed in 3 high-power microscope fields (100x)
[3]average number of goblet cells in an area of 6 contiguous crypts

**Table S6: Primers used for RT-qPCR Analysis**

| Target | Forward Primer | Reverse Primer | Reference |
|---|---|---|---|
| 18S | CATTCGAACGTCTGCCCTAT | CCTGCTGCCTTCCTTGGA | Lab designed |
| Reg3β | ATAGGGCAACTTCACCTCAC | CTGCCTTAGACCGTGCTTTC | Steltler et al. (2011) (3) |
| DEFB14 | GCCTCTTCCAGGTGTTTTTG | GAGACCACAGGTGCCAATTT | Jatana et al. (2008) (4) |
| mCRAMP | GTCTTGGGAACCATGCAGTT | TGGTTGAAGTCATCCACAGC | Park et al. (2011) (5) |

#### Supplemental Figure Legends

**Figure S1: Selection of NCAPD3-targeting shRNA and success rate of NCAPD3KD-targeting *in vivo*. A.** Selection of mouse NCAPD3 targeting shRNA sequence. B6 immortalized mouse macrophage lines lentivirally transduced with 3 different IPTG-inducible mouse NCAPD3 targeting shRNA sequences (1-3) or a non-targeting control shRNA (C). Expression of shRNAs was induced by the addition of 2 mM IPTG to the culture medium daily for 2 days. Cell lysates analyzed by immunoblot for NCAPD3 expression relative to Na/K ATPase loading control. Construct #3 was chosen for inclusion in the mouse transgene targeting cassette. **B.** Success rate of transgenic mouse creation. Three injection conditions with different ratios of Cas9/sgRNA/targeting vector were used to create targeted B6SJL/F2 fertilized oocytes that were then transferred to CD1 pseudo-pregnant recipients. Resulting pups (Founders) were then screened by qPCR and confirmed by sequencing of PCR products for transgene insertion and orientation. Of the mice with correctly oriented transgenes, germline transmission occurred in 40% of the lines. Three lines were crossed to villin-rtTA mice for experimental use. These lines were backcrossed to a C57BL/6 background for 5-7 generations prior to experimental use.

**Figure S2: Flow cytometry gating for intestinal epithelial NCAPD3 protein expression analysis. A.** Analysis of intestinal cells from driverless NCAPD3 KD mice treated with 9-TB-dox (25 mg/kg i.p.) for 4 days (control colon). **B.** Analysis of intestinal cells from NCAPD3 KD mice treated with 9-TB-dox (25 mg/kg i.p.) for 4 days.

**Figure S3: Cytokine and chemokine profile of DSS-treated NCAPD3 KD mice.** Endpoint colonic tissue lysates from colitic NCAPD3KD mice treated with saline or 9-TB-dox were assayed using a 19-plex MSD panel that included: IL-1β, IL-2, IL-4, IL-5, IL-6, IL-9, IL-10, IL-12p70, IL-15, IL-17AF, IL27p28/IL-30, IL-33, IFNγ, IP10, KC-GRO, MCP1, MIP1α, MIP2, and TNFα. All significantly different comparisons are shown. Mean ±SEM graphed; n=9 mice/group. Significance determined by unpaired 2-tailed t-test. **A.** Tissue levels of proinflammatory chemokines. **B.** Tissue levels of anti-inflammatory cytokines. **C.** Tissue levels of proinflammatory and epithelial barrier-damaging cytokines

**Figure S4: Impact of 9-TB-Dox on *Salmonella* viability *in vitro* and *in vivo*. A.** Comparison of antibiotic activity of doxycycline and 9-TB-dox on *Salmonella enterica* serovar Typhimurium ΔAroA survival *in vitro*. Bacteria were cultured with the indicated drug concentration for 20 hours at 37°C and endpoint absorbance graphed. Data are representative of 3 independent experiments. **B-D.** Comparison of *Salmonella* recovery from mice treated with either saline or 9-TB-dox (25 mg/kg i.p. daily) over 7 days from stool (**B**), cecal tissue (**C**), or distal colon tissue (**D**). Tissue loads determined on day 7 post-infection. Mean ±SEM; n=2 female mice/group

**Figure S5: Raw and uncropped immunoblot images. A.** NCAPD3 and tubulin immunoblots displayed in Figure 1. **B.** NCAPD3 and Na/K ATPase immunoblots displayed in Supplemental Figure 1. Immunoblots were sequentially probed with NCAPD3 then loading control antibodies. Bands of correct molecular weight for target indicated by arrows.

**A**

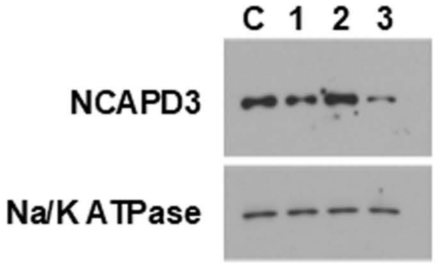


**B**

| Injection ID | Founders | Transgene Positive | Correct Orientation |
|---|---|---|---|
| 20p/20/2 | 5 | 0 | 0 |
| 5p/5/0/5-1 | 42 | 14 | 12 |
| 5p/5/0.5-2 | 31 | 6 | 5 |
| Total Mice (% of Founders) | 78 | 20 (25.6%) | 17 (21.6%) |

**Figure S1**

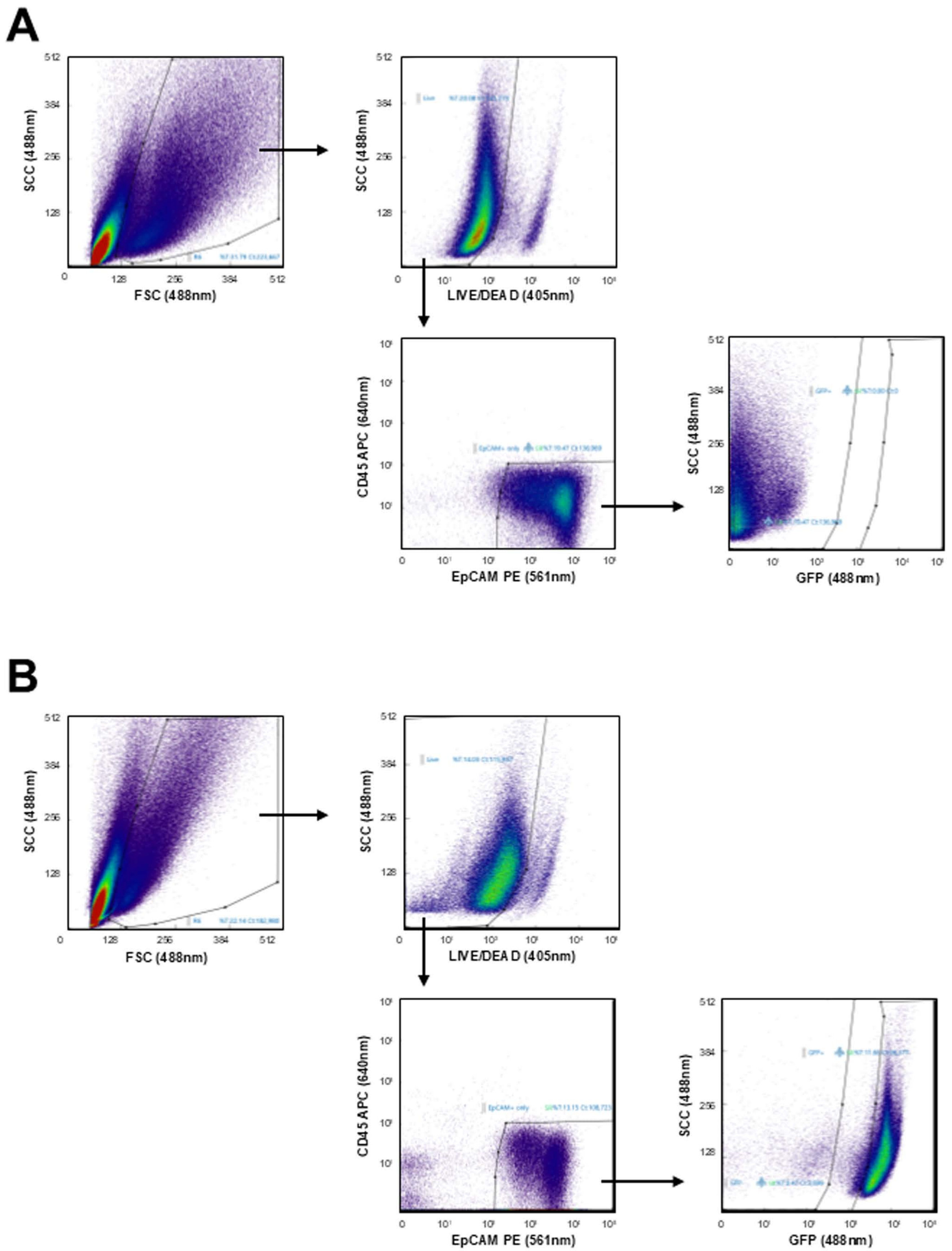
A
SCC (488nm)
FSC (488nm)
LIVE/DEAD (405nm)
CD45 APC (640nm)
EpCAM PE (561nm)
GFP (488nm)
B

Figure S2

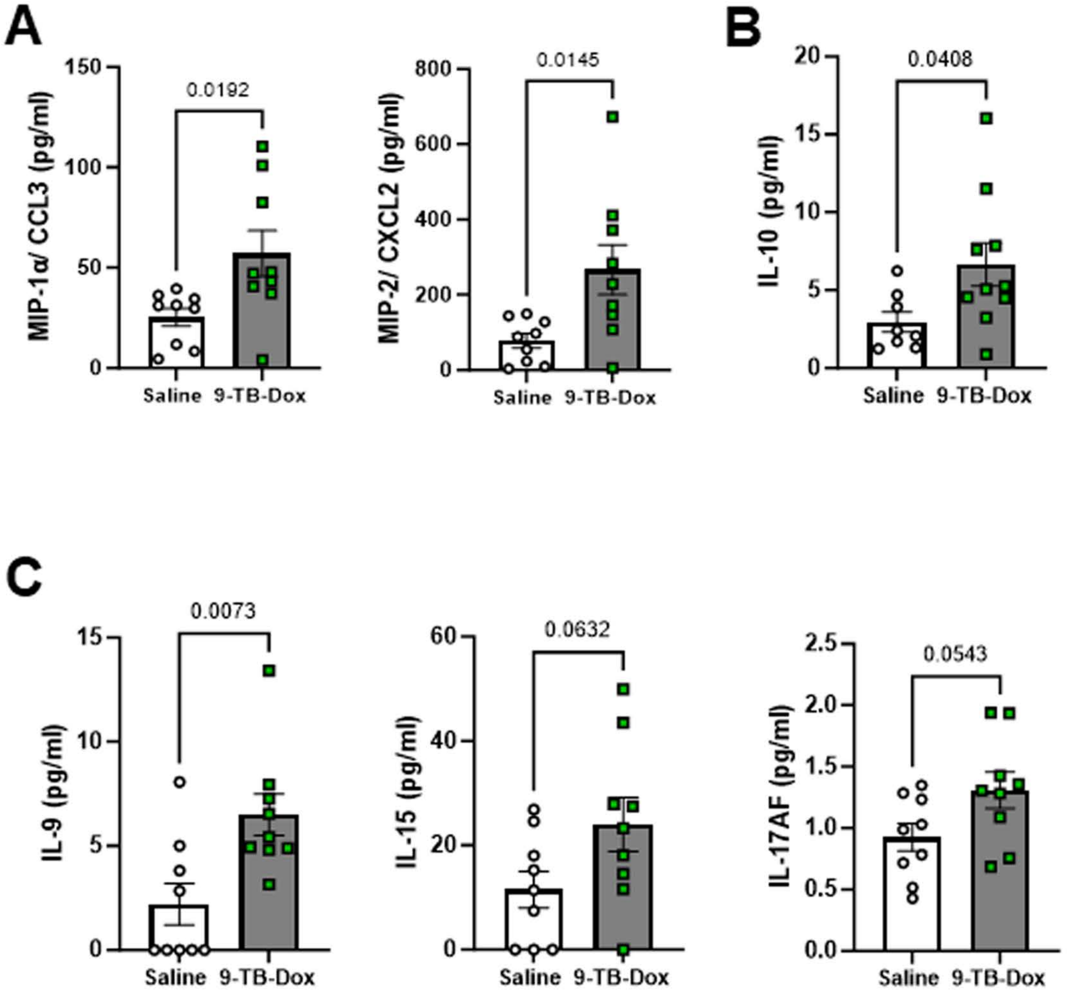
A
MIP-1α/ CCL3 (pg/ml)
150
100
50
0
0.0192
Saline
9-TB-Dox
MIP-2/ CXCL2 (pg/ml)
800
600
400
200
0
0.0145
Saline
9-TB-Dox
B
IL-10 (pg/ml)
20
15
10
5
0
0.0408
Saline
9-TB-Dox
C
IL-9 (pg/ml)
15
10
5
0
0.0073
Saline
9-TB-Dox
IL-15 (pg/ml)
60
40
20
0
0.0632
Saline
9-TB-Dox
IL-17AF (pg/ml)
2.5
2.0
1.5
1.0
0.5
0.0
0.0543
Saline
9-TB-Dox

### Figure S3

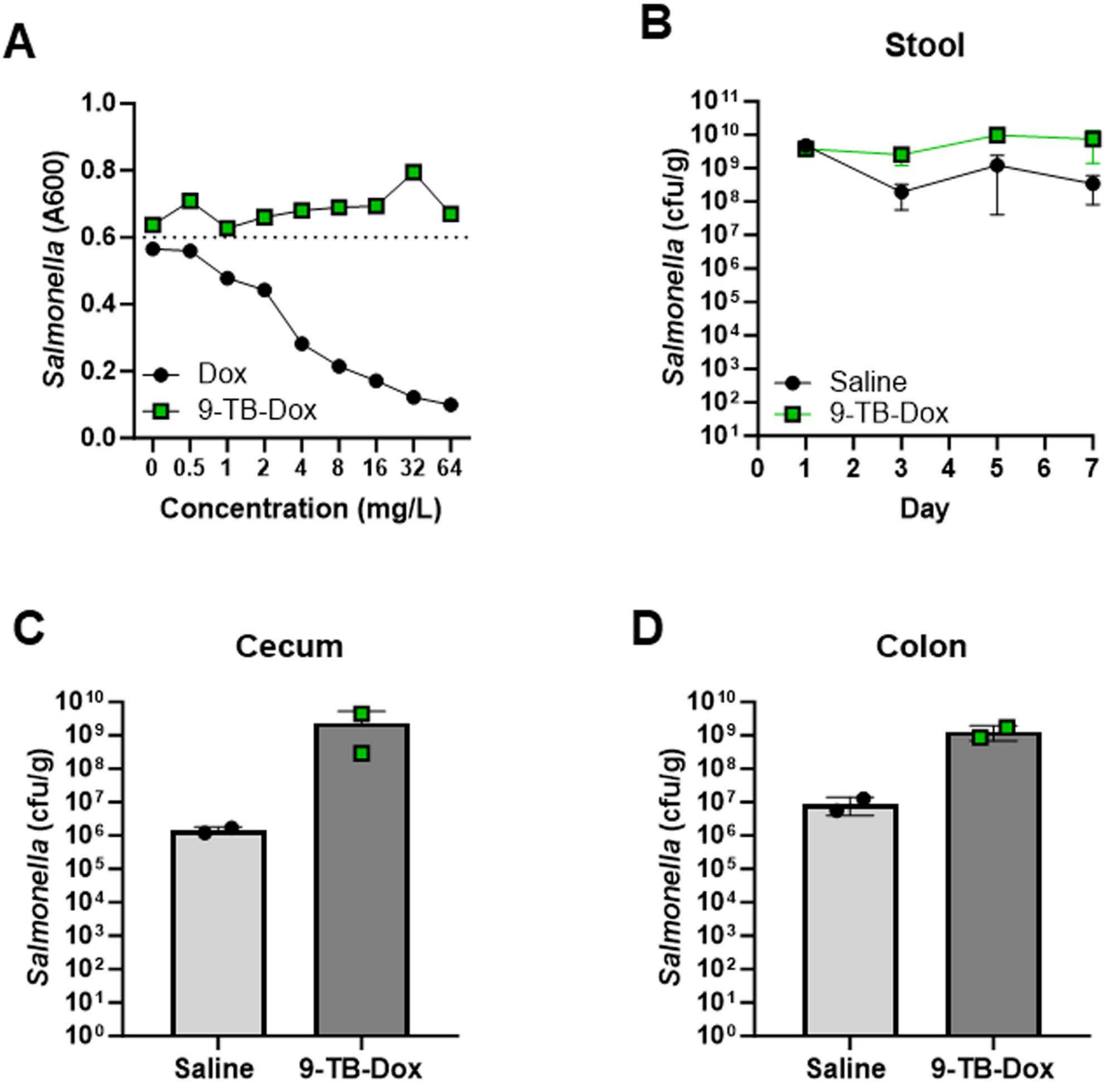
A
1.0
0.8
0.6
0.4
0.2
0.0
Salmonella (A600)
Dox
9-TB-Dox
0
0.5
1
2
4
8
16
32
64
Concentration (mg/L)
B
Stool
$10^{11}$
$10^{10}$
$10^{9}$
$10^{8}$
$10^{7}$
$10^{6}$
$10^{5}$
$10^{4}$
$10^{3}$
$10^{2}$
$10^{1}$
Salmonella (cfu/g)
Saline
9-TB-Dox
0
1
2
3
4
5
6
7
Day
C
Cecum
$10^{10}$
$10^{9}$
$10^{8}$
$10^{7}$
$10^{6}$
$10^{5}$
$10^{4}$
$10^{3}$
$10^{2}$
$10^{1}$
$10^{0}$
Salmonella (cfu/g)
Saline
9-TB-Dox
D
Colon
$10^{10}$
$10^{9}$
$10^{8}$
$10^{7}$
$10^{6}$
$10^{5}$
$10^{4}$
$10^{3}$
$10^{2}$
$10^{1}$
$10^{0}$
Salmonella (cfu/g)
Saline
9-TB-Dox

**Figure S4**

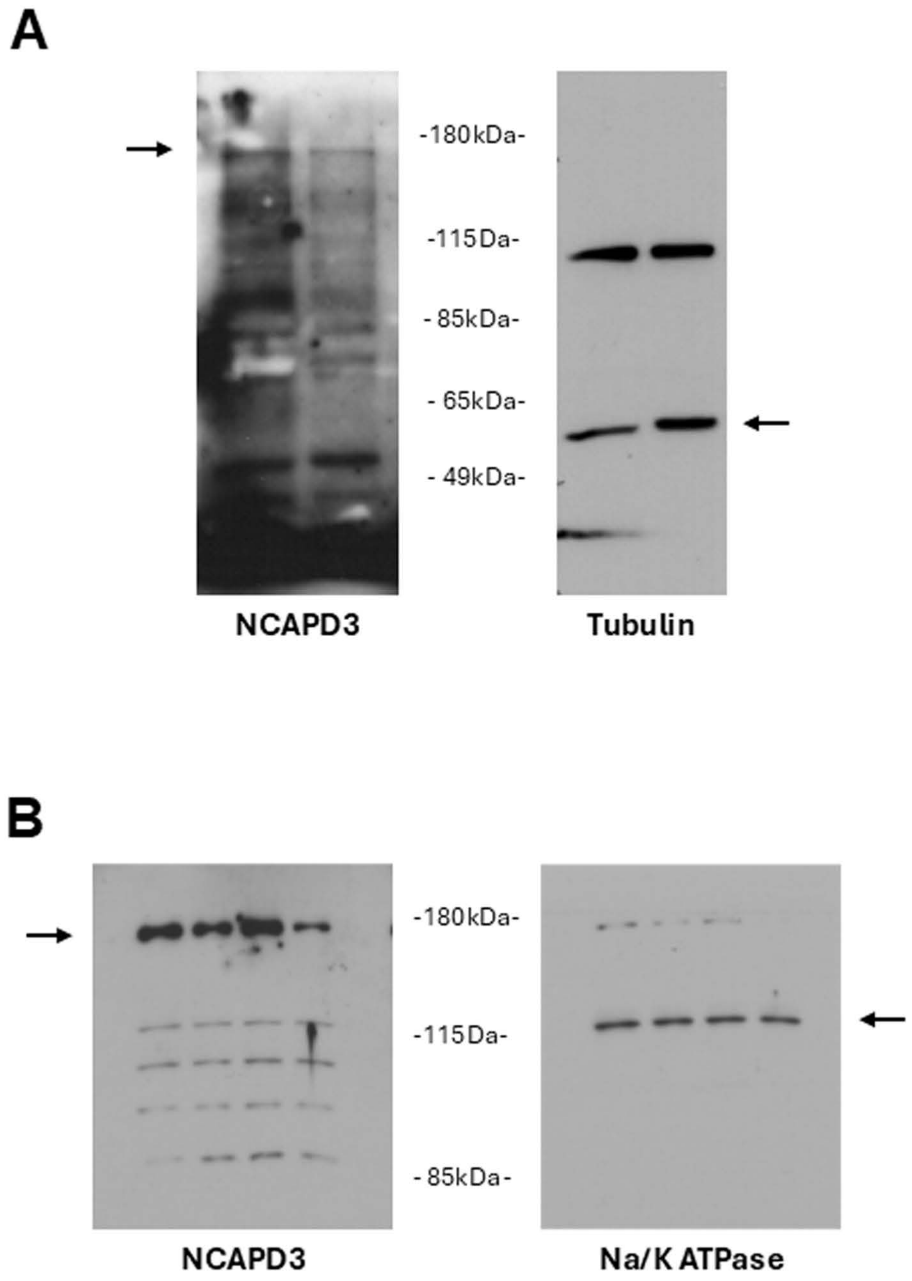
A
-180kDa-
-115Da-
- 85kDa-
- 65kDa-
- 49kDa-
NCAPD3
Tubulin
B
-180kDa-
-115Da-
- 85kDa-
NCAPD3
Na/K ATPase

**Figure S5**
